# Biofilm Formation and Niche Adaptation Play Fundamental Role in Shaping Long-Distance Transported Dust Microbial Community

**DOI:** 10.1101/2024.11.14.623531

**Authors:** Naama Lang-Yona, Hilit Levy Barazany, Talal Salti, Rahamim Ben Navi, Liat Rahamin Ben-Navi, Ilana Kolodkin-Gal

## Abstract

The long-distance transport of microorganisms via dust events is expected to become more prominent and intensified. At the same time, the survival mechanisms of air/dust-borne bacteria and their possible contribution to global processes remain poorly understood. In this study, we characterized *Bacillus* species from transitional season dust storms, previously identified as a significant component of the bioactive community of the dust microbiome. Our results demonstrated substantial growth and biofilm formation diversification, potentially linked to niche adaptation and surface-associated biofilm formation within heterogeneous dust particles. Most dust isolates form biofilms while exhibiting different preferences for media composition. Sterile dust induced biofilm formation, growth, matrix gene expression of *B. subtilis*, and robust biofilms in key related dust isolates. Overall, our results highlight the significance of biofilm formation as an adaptive mechanism in the distinct habitat of dust storms, with niche adaptation potentially playing a role in microenvironments within the dust particle. These results hold significant potential for implications on terrestrial and aquatic ecology and health, suggesting a pivotal process by which bacteria survive and evolve in this understudied habitat.

## Introduction

Bioaerosols, also referred as primary biological aerosols, include a diverse group of microbes directly released from the biosphere into the atmosphere (1-5). Bioaerosols can significantly impact terrestrial (6) and aquatic ecosystems (7), affect biogeochemical (8) and hydrological cycles (9), and impact human health (6, 10).

Dust events have been associated with health hazardous effects (11, 12) and identified as significant aerial carriers of microorganisms (13, 14), playing an important role in the long-distance dispersion of bioaerosols (15-17), including potential plant and human pathogens (18). However, the associated risks, ecological and biogeochemical impacts are still largely unknown (19). The chemical and physical properties of dust particles may influence their interaction with biological matter, affecting bacterial transport (20), virulence (21, 22), and biofilm formation (22, 23). The projected intensification of atmospheric dust loadings due to desertification and land-use changes (24, 25) highlights the urgent need for a better understanding of the impact of dust particles as well as their microbiome (19). A comprehensive understanding of the dust microbiome community composition is commonly obtained through metagenomic studies (26-29). However, to gain deeper insights on the survival and physiological processes of key members of the dust-borne community, follow up experimental setups are required.

The potentially active dust-borne community previously identified by us (26) and others (29) included Firmicutes (also known as *Bacillota*), many of which are spore formers. Specifically, the *Bacillus* genus has been shown to promote biofilm formation in the presence of clay minerals (23). This ability suggests that biofilm formation, rather than sporulation, holds a potential advantage for *Bacillus* within the airborne dust environment. The dust particulates constitute an extreme and unique ecosystem.

During dust storms, dust-*Bacilli* must attach to dust particles and survive with minimal nutrients while vectored to new, unknown niches for potential colonization. In a previous study, we explored the bioactive dust-borne bacterial community and their associations through the 16S rRNA gene and transcriptional signatures of the sampled dust (26). The Firmicutes phylum, with a predominance of *Bacillus* (26), showed the highest RNA/DNA ratio, indicating a high potential for shaping dust microbiome. Nevertheless, to date, it is not known if such spore-forming dust-borne bacteria are dormant during transport or if they harbor vegetative form aboard the dust particles. In this study, we isolated bacteria from the same dust samples that were previously characterized for their bioactive microbiome (26). We analyzed the characteristics of growth and biofilm formation in multiple isolates and their relevance to biofilm formation on dust. Collectively, our results suggest an appealing hypothesis regarding the contribution of biofilm formation to the fitness of air-borne bacteria in this underexplored habitat.

## Results

To confirm the presence of *Bacilli* in the dust samples, as predicted in our previous study, and allow subsequent analysis (26), we enriched spore formers from collected dust samples during both spring and fall dust samples (See **Fig 1A** and **Table S1**), as conducted previously from soil (30), harvesting *Bacillus* isolates. These isolates included but were not limited to a broad repertoire of *Bacillus* species, e.g., Is_2H - *Bacillus* sp. 98.36% identification for GyrA gene); Is_2J - *Bacillus mojavensi*s (92.95% for GyrA gene); Is_2R-*Priestia megaterium* (98.75% for GyrA gene), Is_1.4 -*Metabacillus* sp. (82.27% for GyrA gene), Is_1.28-*Bacillus altitudinis* (97.76% for GyrA gene) and Is_2.8 *Bacillus licheniformis* (89.05% for GyrA gene) were further identified genomically through sanger sequencing based on 16S rRNA and GyrA for *Bacillus* (**Fig. 1B & C; Table S2 & S3**) (31). As shown, phylogenetic distances present different *Bacilli* strains on dust in both sampled seasons (14 strains for the 16S rRNA gene and 10 strains for the GyrA gene with genomic distance > 0.05). Genomic distances appear to be bigger for the 16S rRNA gene, most probably due to the larger size of this amplicon (See **Table S3**). In addition, it seems that genomic distances are not always consistent between the two genes (e.g., for Is_2H, Is_1.4, Is_2B, Is_2J) indicating possible new *Bacillus* strains necessitating shotgun sequencing of whole genome identification.

Once we characterized *B. subtilis* and related spore formers residing in dust, we conducted a study on biofilm formation and growth in three different environments: a defined buffered medium, MSgg, known for inducing highly structured biofilms in *B. subtilis* (32), as well as in the rich B4 medium with and without a calcium source. These media have been shown to induce biofilm formation in *B. subtilis, M. smegmatis*, and other *Bacillus* species (32-34).

**Figure 1.**
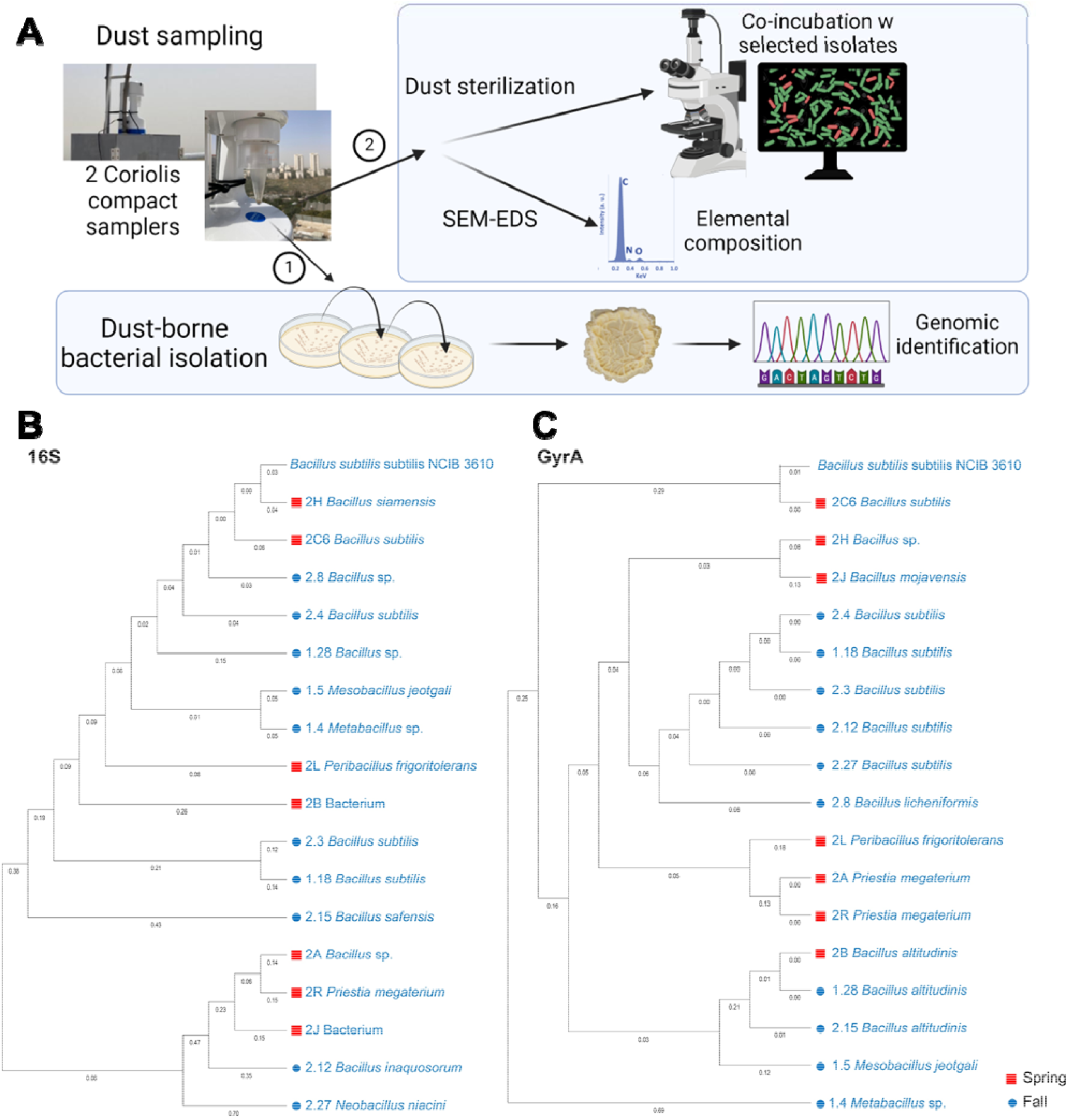
Dust *Bacillus* isolates demonstrate genomic diversity. Experimental design of the study (**A**), and the Maximum likelihood phylogenetic trees for the 16S rRNA (**B**) and the GyrA (**C**) genes, calculated for the paired end sanger sequenced isolates from both spring (red squares) and Fall (blue circles) dust samples. Distances smaller than 0.05 were removed for clarity. Experimental design prepared using *BioRender.com*

The growth and carrying capacity in isolates from both events were diverse (**Fig. S1A & B**). Generally, dust isolates showed optimal growth in the defined medium MSgg (**Fig. S1C**).

This intriguing finding suggests a strong compatibility between the defined, nutrient-limited medium and the dust habitat, shedding light on the adaptability of these isolates.

Is_2H, Is_1.4, Is_2B, and Is 2J, which represent diverse *Bacillus* species, exhibited varying growth rates and maximum optical densities under different growth conditions. Similar patterns were observed with Is_2R and Is_2A from the *Prisela megaterium* genus.

The ability to form biofilm was shown to be beneficial for environmental microorganisms for survival and communication (35). We hypothesized that biofilm formation may be essential for surviving airborne dust particles and may not be a direct readout of the growth parameters. To test this, we first assessed the formation of complex colony biofilms of each isolate across the different media.

Our research findings revealed intricate 3D pattern formation in all isolates, as depicted in **Fig. 2**. Notably, specific *B. subtilis* and non-B. *subtilis* isolates like Is_2J, Is 2B, Is_1.28, and Is_1.5 showcased highly intricate biofilms across all growth media (**Fig. 2A & B**). On the other hand, isolates such as Is_2R and Is_2.27 displayed distinct pattern formations on select media types, indicating diverse behaviors among the isolates. Largely all isolates generated complex morphology under at least one condition hinting at a potential necessity for secreted extracellular polymers.

**Figure 2.**
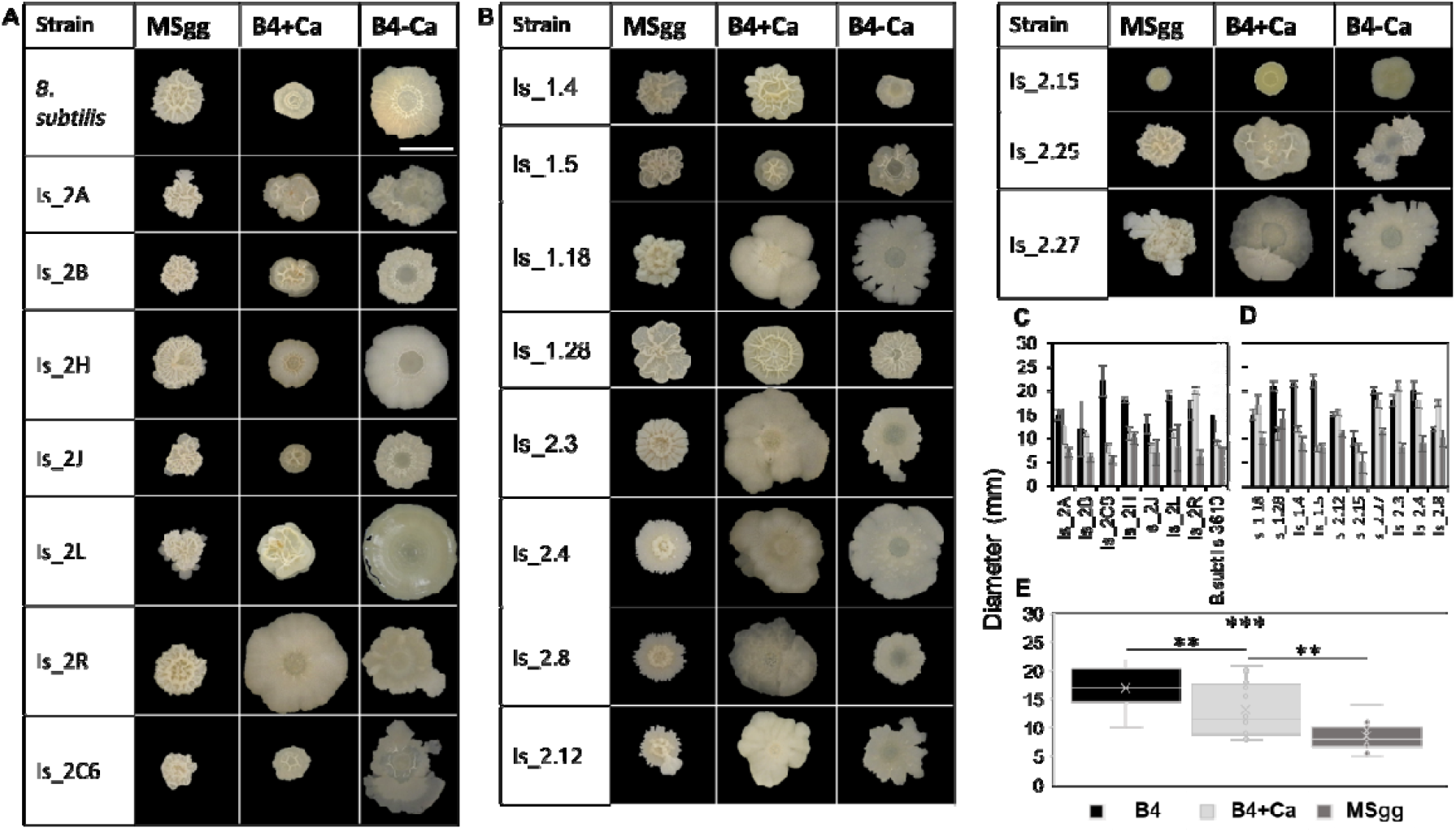
Biofilm formation by dust *Bacilli*. Morphological structures of spring (**A**) and fall (**B**) bacillus isolate three types of media: MSgg defined medium, B4 enriched with calcium (B4+Ca) and B4 without calcium (B4). Scale bar = 10 mm. The diameter of the colonies at the different media are shown for spring (**C**) and fall (**D**) dust samples. Two-way ANOVA, with Tukey post-hoc between media, with confidence level = 0.99. Experiments were performed 3 independent times in triplicates.

A quantitative analysis challenged the expectation that more intricate structures would result in smaller diameters as a compensatory response (**Fig. 2C & D**). We found that the average diameter in isolates from both dust events was largest in B4 medium, and significantly smallest in MSgg medium (**Fig. 2E**; ANOVA, with Tukey post-hoc *p* < 0.001). These findings suggest an inverse correlation between complex structures and colony diameter, and prompt further investigation into the underlying mechanisms driving these observations.

Floating biofilm formation was quantified as described in the materials and methods, with more variance observed in B4+Ca compared with MSgg. No pellicle was observed in B4 (**Fig. S2**).

Notably, biofilm formation on solid medium did not predict the success of a given isolate of generating a pellicle in each medium. Moreover, some isolates e.g., Is_2.15 failed to generate complex colonies while excelling in pellicle formation (**Fig 2B & S3**). While incapable of floating biofilms, most isolates formed a submerged biofilm in B4 (**Fig. S3**).

In order to explore the potential connections between growth patterns and biofilm formation, we examined the expansion of biofilm colonies on agar plates. This expansion reflects various factors such as biomass, motility, and to some extent, the production of exopolymers on the surface. We found that the growth rate and maximum optical density (OD) were correlated, especially in B4, suggesting that both are related indicators of planktonic growth (**Fig. 3A**).

**Figure 3.**
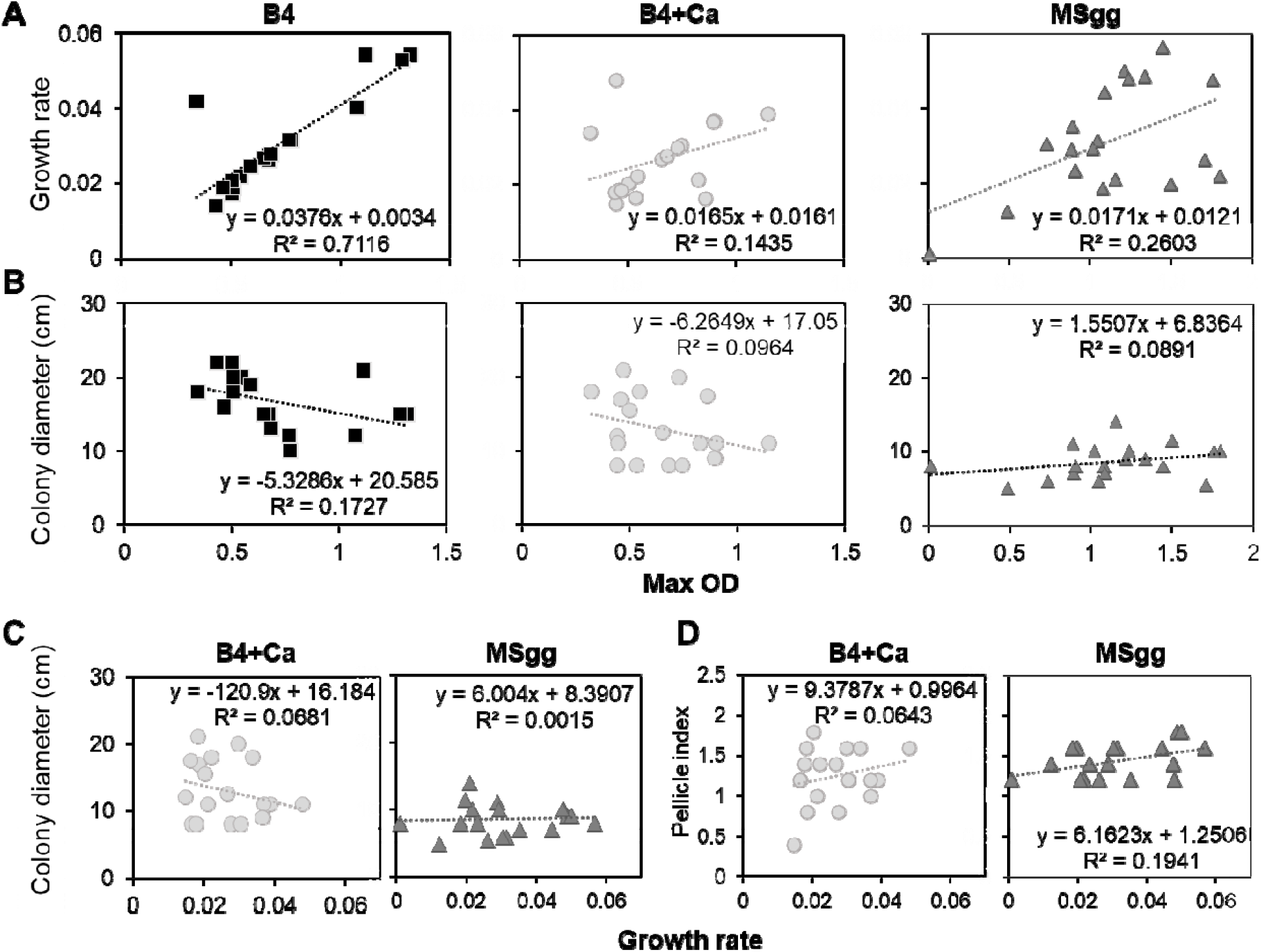
Growth parameters of dust isolates. **A.** planktonic growth rates, and **B.** Colony growth diameter versus the maximum planktonic optical density (OD) for isolates grown up to 72 h at 30°C, comparing B4 medium, B4+Ca and MSgg synthetic medium. **C** Colony growth diameter, and **D** Pellicle complexity index versus planktonic growth rate for isolates grown up to 72 h at 30°C, comparing B4+Ca medium and MSgg synthetic medium.

The size of the colony was not accurately predicted by either the maximum OD or growth rate (see **Fig. 3B & C**). For pellicle formation correlation was more evident (**Fig. 3D**). These findings suggest that the formation of biofilms on a solid surface differs significantly from growth, while the development of floating biofilms in these closely related genomes directly reflects the growth rate.

The dust particles are diverse in terms of their physical and chemical composition (**Fig. S4**). Hence, the ability of dust-isolated bacteria to thrive on artificial media may not necessarily reflect their growth proficiency while being transported over dust particles. To explore the potential of *Bacilli* to proliferate in a dust environment, we further examined their growth and biofilm formation on natural dust particles. This involved incubating sterilized dust samples with a specific genetically manipulatable *Bacillus subtilis* strain (*B. subtilis* 3610).

While the dust provided some of the necessary trace elements and nutrients for microbial growth **(Fig. 4A)**, the application of a single external nitrogen source (glutamine) (38) significantly enhanced biofilm formation on the dust, which had positive effects on cell growth **(Fig. 4B)**. Additionally, a single carbon source, glycerol (39), also improved biofilm formation and growth **(Fig. 4A & B)**, although not to the same extent as glutamine. These results support the primary barrier to the survival and biofilm formation of heterotrophic *B. subtilis* on dust is nitrogen/nitrogen fixation but also, most importantly, that dust particles originated from dust storms support biofilm formation.

**Figure 4.**
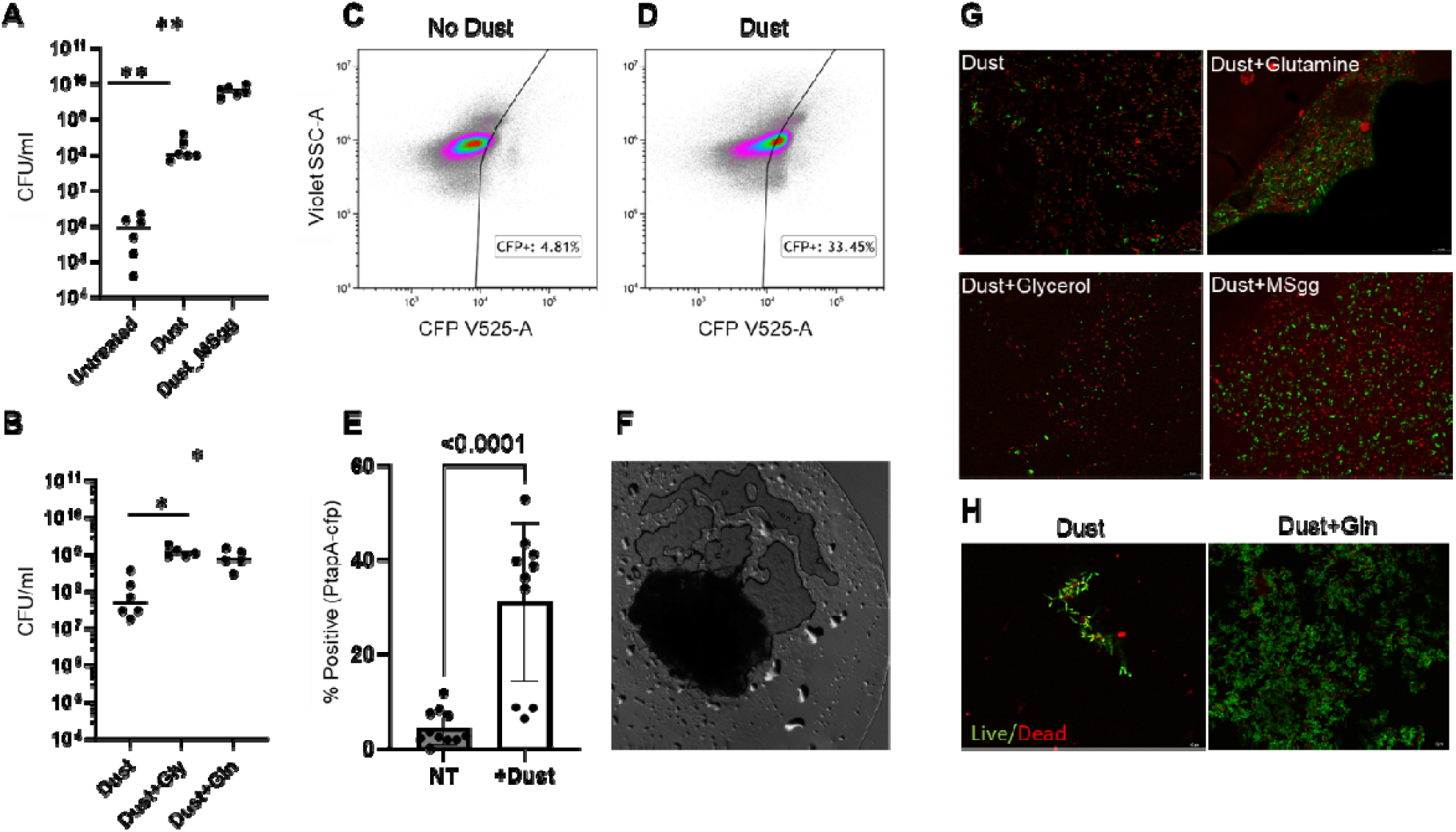
*Bacillus* growth and sporulation abilities over dust. **A.** Colony forming units of *B. subtilis* 3610 grown on either 1.5% agar, Agar applied with sterile dust, and MSgg-Agar. **B.** Colony forming units of *B. subtilis 3610* grown on either Agar applied with sterile dust, agar applied with sterile dust and 0.5% glycerol (Gly) and agar applied with 0.5% glutamine (Gln). **C-D** Expression of the P_tapA-CFP_ either agar or agar applied with sterile dust (n=9). **E.** % of *CFP* positive bacterial cells in the presence and absence of dust/ (D-E) Experiments were performed 9 independent times due to the variability of dust particle**s F.** The magnification (x63) of a film (full arrow) associated with sterile dust particles **G.** *B. subtilis* strain carrying TasA-mcherry, and P_hag_-gfp was grown on dust either applied with Gly or Gln (0.5%) or MSgg (10%) and imaged with confocal microscopy with x63 magnification. **H.** Dust isolate Is_2H grown over sterile dust with and without the addition of glutamine (Gln) and stained with a live/dead stain as described in materials and methods at indicated conditions. The representative fields represent 3 independent experiments performed in duplicates

In previous studies, genes required for the assembly of the complex structures such as the adhesin and putative amyloid protein *TasA* were shown to play an important role in plant and soil colonization (36). We observed additional evidence of the activation of the specific regulation of the biofilm developmental program on dust. We measured the expression of the *B. subtilis tapA-sipW-tasA* promoter tagged to Cyan fluorescent protein (CFP), which was from a single copy in the *B. subtilis* chromosome. Our results showed that the interactions with dust significantly increased (five-fold, *p*-value < 0.0001) the transcription of this promoter in a highly liming growth media (0.5% MSgg) over multiple experiments **(Fig. 4G)**.

This effect was consistently observed despite the measured variability of dust particles (**Fig. S4**). Moreover, *B. subtilis* formed structured biofilms on sterile dust, where cells expressed both *TasA* adhesin tagged with mCherry and the flagellin protein (as judged by the transcriptional fusion P_hag_-GFP **(Fig. 4G)**.

To further confirm biofilm formation on dust with our dust-borne isolates, we selected two isolates, Is_2H (identified as *Bacillus sp*. at the genus level) and Is_2.4 (identified as *B. subtilis*), both showing high ability to form biofilms on three different media (**Fig. 2**). We then incubated these isolates with sterile dust particles as a source of nutrients. Both isolates formed biofilms on the dust, especially when glutamine was present as a source of nitrogen. Is_2H showed independence from glutamine (**Fig. 4H**), while Is_2.4 relied on glutamine (**Fig. S5**). We assessed the viability of the cells using a Live/Dead stain and found that there was a significant presence of viable cells within the dust-associated biofilm population, particularly in the presence of a nitrogen source.

## Discussion

In this study we explored the characteristics of bacteria living in a highly competitive and extreme environment – dust particulates, focusing on *Bacillus* isolates as our case study. By exploring the mechanisms of bacterial biofilm formation and survival on and within dust, we expect to shed light on a poorly characterized aspect of our air microbiome, as potentially modified and regulated by the dust particulate environment. A major advantage of exploring the dust environment is its highly selected and most unique bacterial community, presenting many skills for fitness and survival. It is anticipated that the viable fraction of airborne microbes will possess distinct resistance properties against the extreme conditions within the atmospheric aerosols including desiccation, UV radiation, and other stressors, promoting competition among the dust-borne microorganisms. Nevertheless, although specific bacterial species have been isolated from atmospheric samples (37-39), their unique properties and characteristics have been limitedly explored to date.

While sporulation was frequently considered in this context, biofilm formation where polymers are secreted, can also serve as an efficient protective buffer against environmental stressors. Microbial exopolymers can serve both as antimicrobial barriers and as osmo-protectants (40), and therefore may enable the survival and attachment of bacteria during airborne transport. Consistently, we found that most dust isolates formed robust biofilm under laboratory conditions (**Fig. 2**). Pellicles formation correlated with the growth rate during planktonic growth while complex colony formation did not, potentially indicating two different axes of niche adaptation: one for growth and one for surface-associated pattern formation (**Fig. 3**). Additionally, a well-characterized strain of *B. subtilis* 3610 expressed flagellin and biofilm adhesins robustly on dust (**Fig. 4**). Furthermore, the growth over sterilized dust collected during the dust events strongly induced the transcription of the biofilm associated operon, *tapA-sipW-tasA*, which encodes an accessory protein, a signal peptidase and an amyloid-like adhesin (41), all essential for biofilm formation (**Fig. 5**).

**Figure 5.**
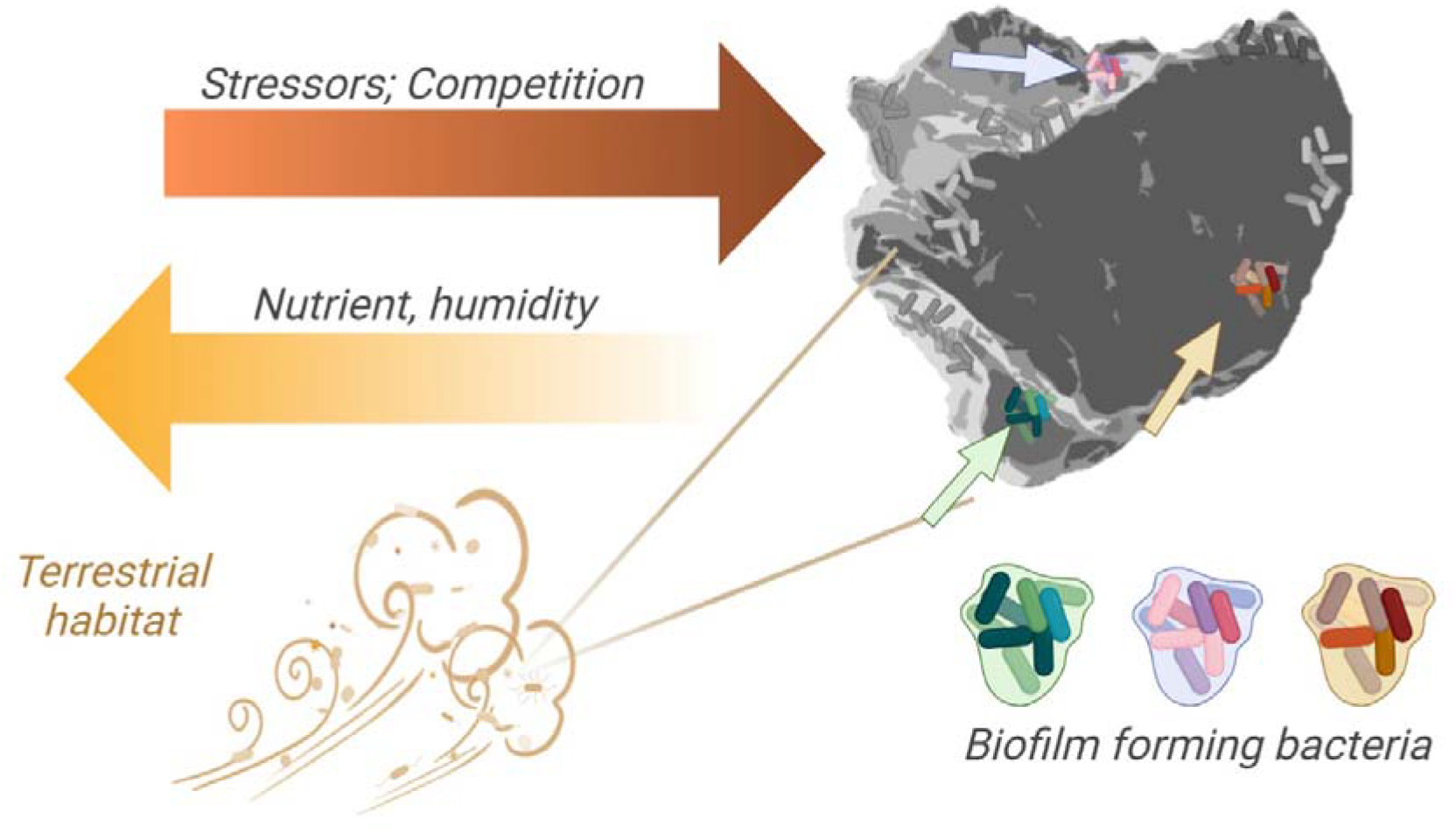
Stressors and competition shape biofilm-forming communities on dust. A conceptual model explaining the diverted microbial communities detected in dust, shaped by environmental conditions and stressors such as reduced humidity, low nutrient availability, radiation. Created using *BioRender.com*

Interestingly, our analysis of these isolated organisms revealed strong evidence for specialized adaptation and reprogramming of the organism’s environmental response networks, including sensors, signal transducers, and transcriptional and post-transcriptional regulators. Even closely related *Bacillus* isolates (**Fig. 1**) were found to form optimal biofilms under different conditions (**Fig. 2**). This finding may suggest that there are multiple microenvironments within the dust, consistent with the heterogeneity in shape and chemical composition of the dust particles **(Fig. S5)**.

Growth on dust is feasible due to efficient bioleaching and the presence of carbon and nitrogen in the dust particles. These nutrients may originate from predation or active metabolism of the dust-borne microorganisms. The addition of external nitrogen or carbon from the environment can improve biofilm formation, suggesting a dependence on microorganisms capable of carbon and/or nitrogen fixation. Further research is needed to understand the symbiotic relationship between different community members.

Our results on biofilm formation abilities of bacteria carried by dust aerosols shed light on how microorganisms may survive the extreme conditions during their atmospheric journey. Unravelling the ecological interactions, chemical communication, and biological regulation of these dust-borne communities holds significant implications for understanding their potential impact on terrestrial and aquatic ecosystems at their destination. Additionally, exploring the connections between biofilm biology and biogeochemical cycles could be highly valuable. Particularly interesting is the carbonate mineralization of airborne *Bacillus* biofilms, and its potential role in carbon dioxide accumulation and sequestration.

## Materials and Methods

### Media

For Fluorescent microscopy, we used *B. subtilis* NCIB 3610 carrying both the P_hag-GFP_ (motility, green) and TasA-mCherry (matrix, red) reporters, lacA::P_hag-gfp_(mls), amyE::P_tapA-tapA-sipW-tasA-mCherry_(cam) (42). For flow cytometry we used a strain carrying ameE:P_tapA-mKate_ (spec) (42). Selective media for cloning purposes were prepared with LB broth. Starter cultures of all strains were prepared using LB (Luria-Bertani) broth (Difco). B4 (0.4% yeast extract, (Difco), 0.5% D-glucose and 0.25% calcium acetate (Sigma Aldrich)) was prepared as described previously. When indicated, strains were grown in MSgg medium [5 mM potassium phosphate, 100 mM MOPS (pH 7), 2 mM MgCl_2_, 50 μM MnCl_2_, 50 μM FeCl_3_ for liquid medium or 125 μM FeCl_3_ for solid medium, 700 μM CaCl2, 1 μM ZnCl2, 2 μM thiamine, 0.5% glycerol, 0.5% glutamate, Threonine (50 μg ml−1), Tryptophan (50 μg ml−1), and Phenyl-Alanine (50 μg ml−1)] (31). MSgg plates [1.5% Bacto Agar (Difco)] were prepared on the day of the experiment and air-dried under a laminar flow hood for 40 min before inoculation.

### Dust sampling

Dust was collected during dust storms in spring and fall 2023 on the rooftop of an eight-story building, at the Technion Institute of Technology, Haifa, Israel (See **Table S1**). In each event, air was collected during daytime for 6 hr using a Coriolis Compact air sampler (air sampling rate of 50 L m^-1^). Before each sampling, the instrument was wiped with 70% ethanol and blank sample was collected by operating the sampler for 30 s in the biological cabinet, to make sure no contaminants are sampled from the instrument itself. Then, sampler inlets were sealed with parafilm and placed on the rooftop for dust sampling into a sterile empty tube.

### Cultivation and isolation of spore formers

After sampling, the Coriolis sample tubes were rehydrated under sterile conditions with 1 ml of sterile phosphate-buffered saline (PBS), and vortexed for 15 min at room temperature. The sample solutions were then incubated at 80°C for 20 m to enrich spore formers, resilient to these high temperatures, flowed by brief vortexing as previously described for soil spore-formers (30). We plated triplicates from each sample on LB agar plates for 48 h incubation at 30°C.

A total of 41 colonies were isolated from the spring sample and 64 from the fall sample. No colonies were observed in the blank samples. Colonies were further isolated three consecutive times to ensure pure cultures were obtained. Followingly, isolates were plated on MSgg and B4+Ca agar for morphological selection. A total of 18 isolates were selected for genomic identification and further studies on growth and biofilm formation. These were grown at 30°C in LB for genomic extraction and storage in 15% glycerol at -80 °C with duplicates.

### Genomic identification

Isolates at late exponential growth phase were palleted and used for DNA extraction for genomic identification through sanger sequencing. DNA from isolated bacteria was extracted using the ZymoBIOMICS DNA kit and protocol (Zymo, Irvine CA). DNA extracts were amplified based on both the 16S rRNA and the GyrA gene (31, 43) (see details on primers in **Table S2**).

PCR amplification conditions included a 20 μl PCR mix including 1X MyTaq mix (Bioline, London, UK), 0.5 μM of each primer, 2 μl DNA extract, and nuclease free PCR-grade water (Sigma Aldrich). The thermal conditions for the 16S rRNA gene included an initial denaturation step at 95 °C for 1 min, 30 cycles of denaturation at 95 °C for 30 s, annealing at 57 °C, for 30 s, and extension at 72 °C, for 1.5 min. A final extension step at 72 °C was added for 10 min. The PCR products were validated on 1.5% agarose gel. The thermal conditions for the GyrA gene included an initial denaturation step at 95 °C for 1 min, 30 cycles of denaturation at 95 °C for 30 s, annealing at 50 °C, for 30 s, and extension at 72 °C, for 40 s. A final extension step at 72 °C was added for 7 min. The PCR products were validated on 1.5% agarose gel. Amplicons were paired end sequenced using the 3500xl genetic analyzer (Life Technology) and manually assembled using the BioEdit Sequence Alignment Editor software. The assembled and aligned sequences were identified using the BLAST tool from NCBI. Sequence alignment and phylogenetic tree construction was performed using the molecular evolutionary genetic analysis (MEGA) software (v. 11.0.13). The sequences were deposited in the GenBank at NCBI (Project accession numbers: SUB14624732 and 2853878).

### Planktonic growth rate

The growth rate of the isolated *Bacilli* was tested using the microplate reader (Synergy H1 Hybrid Multi-Mode Reader). In general, a single colony from each isolate was incubated in 5 ml LB at 30 °C and 120 RPM shaking until reached the logarithmic growth stage. 1 μl from each culture was added to a 100 μl liquid medium with 8 replica per isolate in a 96 well-plate. Each plate included 8 wells of no-isolate blank. Growth was evaluated in MSgg, B4+ Ca and B4 media (32-34). Plates were grown in 30 °C and 120 RPM shaking, enriched with CO_2_ and OD was measured at t = 0, 4, 8, 24, 48, and 72 h.

### Biofilm essay

Cells from a single colony isolated on LB plates were grown to mid-logarithmic phase in a 3-ml LB broth culture (4 hours at 37°C with shaking). Then, a drop was spotted on solid MSgg medium. For engulfment experiments, colonies were inoculated next to a disc placed at distance of 0.3 or 0.5 cm. Plates were incubated at 30°C for the period indicated in the legend for each figure. For floating biofilms (pellicles), cells from mid-logarithmic phase were diluted 1:100 into 3 ml of liquid MSgg and grown in 12-well plates, at 30°C, without shaking, for 48 hours. Plates were incubated with enrichment of CO_2_ at 30 °C for 72 h. Images of colonies and pellicles were obtained with Nikon Coolpix p950 high resolution camera with scale, and the diameter of colonies was calculated with Image J. (32-34).

### Crystal violat biofilm evaluation

Crystal violet was performed at B4 medium as described previously (ref). Biofilms were grown in 12 well plates for 48 hours at 30°C

### Flow Cytometry

*B. subtilis* 3610::*amyE:P*_*tapA*_*-CFP* cells were harvested after 48 hours from Msgg Agar plates with or without dust particles. For flow cytometry analysis, cells were mechanically separated, suspended in PBS and measured on a CytoFLEX S (Beckman Coulter Life Sciences) by recording 100000 events for CFP fluorescence (405 nm laser excitation coupled with a 525 filter) plotted versus the 405 nm (Violet) laser flourescence. Each sample was analyzed using Kaluza Analysis Software (Beckman Coulter Life Sciences).

### Confocal Microscopy

*B. subtilis* 3610: *lacA::P*_*hag-gfp*_(mls), *amyE::P*_*tapA-tapA-sipW-tasA- mCherry*_(cam) were cultured in 24-well Agar plates with dust particles and with or without MSgg, L-glutamine (specific concentration), and Glycerol (0.5%) for 48 hours under static conditions at 30 °C. After 48 hours, the dust particles and the biofilm were removed and visualized and photographed using a confocal microscope (Mica, Leica). For fluorescence-based identification of live and dead bacteria in the biofilms. Bacteria (Is_2H or Is_2.4 isolates) were cultured in 24-well Agar plates with dust particles and with or without L-Glutamine for 48 hours. After 48 hours, the wells were supplemented with the Live/Dead stain (Filmtracer™ LIVE/DEAD™ Biofilm Viability Kit, Invitrogen) using 3 μl each of SYTO®9 (Green fluorescent) and propidium iodide (PI, Red fluorescent) per ml for 20 minutes. The bacteria were visualized and photographed using a confocal microscope (Mica, Leica).

### Statistical analyses

Multiple comparison tests were performed using ANOVA test, and the statistical analysis was performed with GraphPad software. When appropriate, a paired student *t*-test was used to compare between samples.

## Acknowledgment

The study was funded by internal grants to NLY and IKG. We thank Shaked Farhi, Or Argaman for support in dust sampling and technical aid, Tamar Yona and Yuval Kolodkin-Gal for transferring the materials between campuses. Avihai Nahami for the assistance in the analysis of pellicle formation.

## Data Availability

The data generated or analysed during this study are included in this published article and its Supplementary Information files. The assembled and aligned sequences of the isolated airborne bacteria were deposited in the GenBank at NCBI (Project accession numbers: SUB14624732 and 2853878).

