## Supporting Information for "Biofilm Formation and Niche Adaptation Play Fundamental Role in Shaping Long-Distance Transported Dust Microbial Community"

### Table of content

**Tables**

Table S1. Dust sampling metadata………………………………………………………..……………………….. 2
Table S2. Primers used for sanger sequencing identification……………………………. 3
Table S3. Genomic identification scores for dust isolates……………………………….. 4

**Figures**

Figure S1. Bacillus isolates growth curves………………………………...………… 3
Figure S2 – Pellicle in *B. subtilis* isolates……………………………………………. 4
Figure S3- Pellicle and submerged biofilm quantification……………………………… 5
Figure S4. Elemental composition of dust samples…………………………………..… 6
Figure S5. Dust isolate Is_2.4 grown over sterile dust………………………………..… 7

**References**…………………………………………………………………………… 8

**Table S1. Dust sampling metadata.** The date of sampling, sampling times and duration, the air volume sampled, and spore-forming OTU detected.

| Sample ID | Date | Sampling times | Sampling duration | Air volume | Total spore-forming OTU[1] |
|---|---|---|---|---|---|
| | | UTC+2 | hh:mm:ss | $m^3$ | |
| Spring blank | 25-April-2022 | 11:45 | 00:00:30 | 0.025 | 0 |
| Spring sample | 25-April-2022 | 12:00 - 18:00 | 06:00:00 | 18 | 41 |
| Fall blank | 22-October-2023 | 11:45 | 00:00:30 | 0.025 | 0 |
| Fall sample | 22-October-2023 | 12:00 - 18:00 | 06:00:00 | 18 | 64 |

[1]Growth over LB agar

**Table S2.** Primers used for sanger sequencing identification.

| PCR product | Primer name | Oligonucleotide sequence | Reference |
|---|---|---|---|
| 16S rRNA gene | 16S-27F | 5'-AGAGTTTGATCMTGGCTCAG | (1) |
| | 16S-1492R | 5'-CGGTTACCTTGTTACGACTT | |
| GyrA gene | GyrA3-F | 5'-GCDGCHGCNATGCGTTAYAC | (2) |
| | GyrA3-R | 5'-ACAAGMTCWGCKATTTTTTC | |

**Table S3.** Genomic identification scores for dust isolates.[1]

| Amplicon | Isolate ID | Scientific name | Identity (%)[2] |
|---|---|---|---|
| 16S rRNA gene | Is_2A | *Bacillus* sp. | 98.55% |
| | Is_2B | Bacterium | 97.97% |
| | Is_2H | *Bacillus siamensis* | 95.34% |
| | Is_2J | Bacterium | 97.01% |
| | Is_2L | *Peribacillus frigoritolerans* | 95.61% |
| | Is_2R | *Priestia megaterium* | 75.78% |
| | Is_2C6 | *Bacillus subtilis* | 93.95% |
| | Is_1.4 | *Metabacillus* sp. | 96.37% |
| | Is_1.5 | *Mesobacillus jeotgali* | 96.50% |
| | IS_1.18 | *Bacillus subtilis* | 82.06% |
| | Is_1.28 | *Bacillus* sp. | 90.84% |
| | Is_2.3 | *Bacillus subtilis* | 81.50% |
| | Is_2.4 | *Bacillus subtilis* | 91.24% |
| | Is_2.8 | *Bacillus* sp. | 95.86% |
| | Is_2.12 | *Bacillus inaquosorum* | 79.21% |
| | Is_2.15 | *Bacillus safensis* | 89.01% |
| | Is_2.27 | *Neobacillus niacini* | 91.57% |
| GyrA gene | Is_2A | *Priestia megaterium* | 98.57% |
| | Is_2B | *Bacillus altitudinis* | 97.77% |
| | Is_2H | *Bacillus* sp. | 98.36% |
| | Is_2J | *Bacillus mojavensis* | 92.95% |
| | Is_2L | *Peribacillus frigoritolerans* | 84.73% |
| | Is_2R | *Priestia megaterium* | 98.16% |
| | Is_2C6 | *Bacillus subtilis* | 99.12% |
| | Is_1.4 | *Metabacillus sp* | 96.37% |
| | Is_1.5 | *Mesobacillus jeotgali* | 88.15% |
| | IS_1.18 | *Bacillus subtilis* | 97.36% |
| | Is_1.28 | *Bacillus altitudinis* | 97.76% |
| | Is_2.3 | *Bacillus subtilis* | 97.15% |
| | Is_2.4 | *Bacillus subtilis* | 97.84% |
| | Is_2.8 | *Bacillus licheniformis* | 89.05% |
| | Is_2.12 | *Bacillus subtilis* | 96.75% |
| | Is_2.15 | *Bacillus altitudinis* | 93.07% |
| | Is_2.27 | *Bacillus subtilis* | 94.93% |

[1]Blast parameters: database: core nucleotide database; Organism: bacteria; Exclude: uncultured/environmental sample sequences; Optimize for: Highly similar sequences; Blast identification date: 21-July-2024.

[2]If sequence was given more than one highest score identification, the identity was reduced to lowest agreed identification.

**Figure S1**

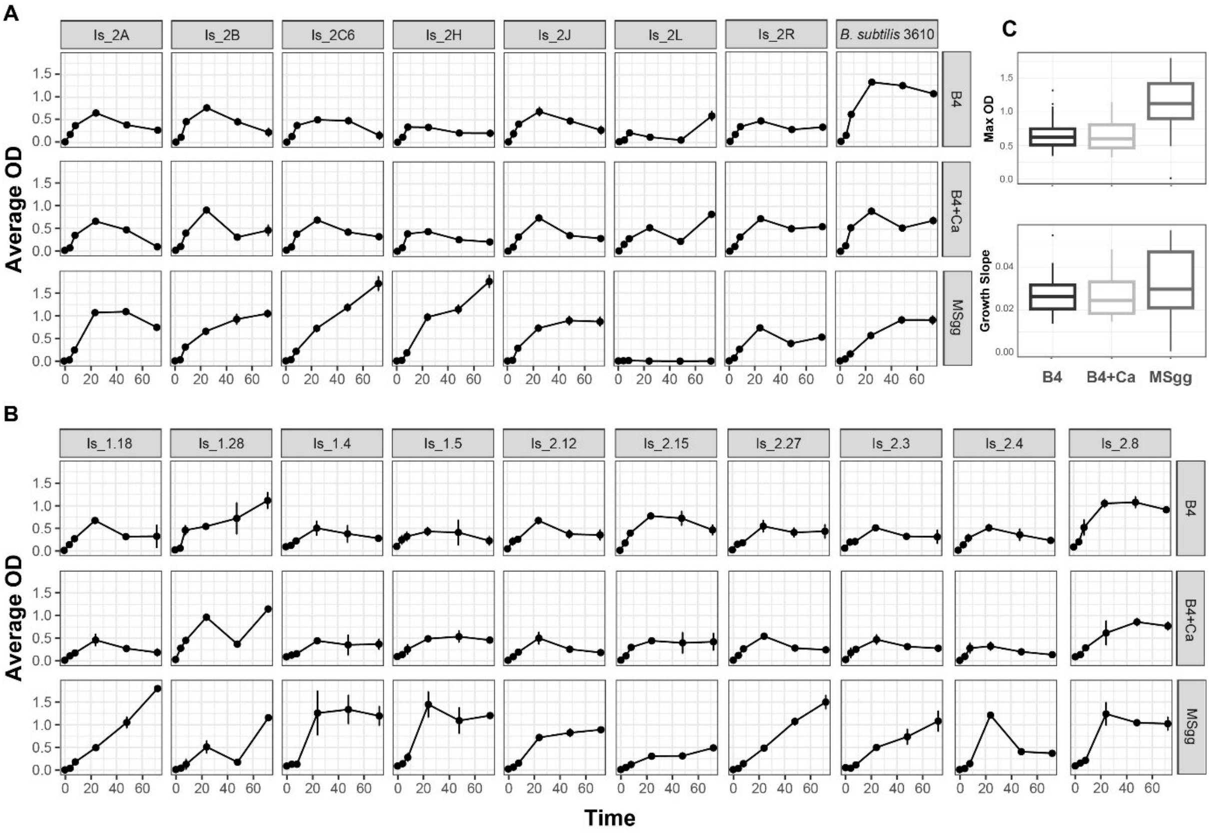


**Figure S1.** ***Bacillus*** **isolates growth curves.** Isolates from spring (**A**) and fall (**B**) samples were monitored for growth rates in liquid media: a defined buffered medium, MSgg, and rich B4 medium with and without a calcium source. The optical density (OD) values represent the average of six replicates after subtraction of the blank OD (medium with no isolate). The maximal OD representing the carrying capacity and the growth slope (**C**), representing the growth rates are presented per each medium. Experiments were performed 3 independent times in triplicates

**Figure S2**

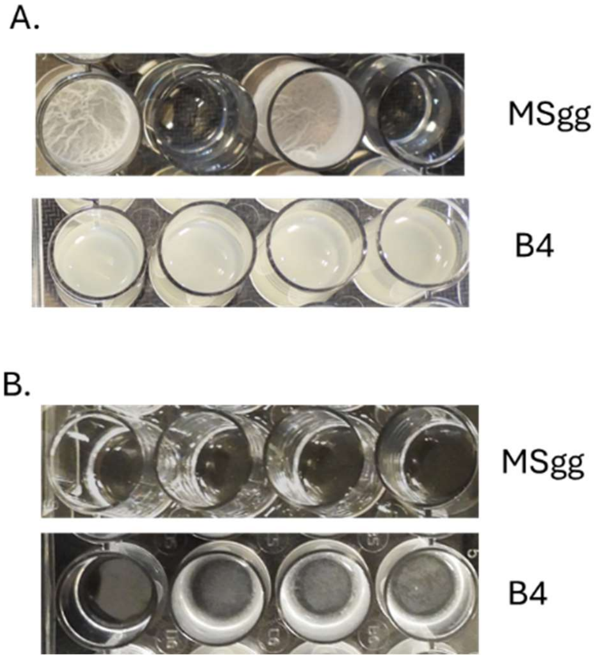


**Figure S2.** ***Bacillus subtilis*** **isolates biofilm formation in MSgg versus B4.** *B. subtilis isolates: NCIB3620,* Is_2H, and Is_2.27, and Is_2.4 (Pellicles were grown in liquid media: a defined buffered medium, MSgg, and rich B4 medium. A. Pellicle images B. Images of the bottom of the well following cells' removal from untreated samples images in A- MSgg – no cellular deposit on the bottom of the well. B4 clear biofilm on the bottom of the well

**Figure S3**

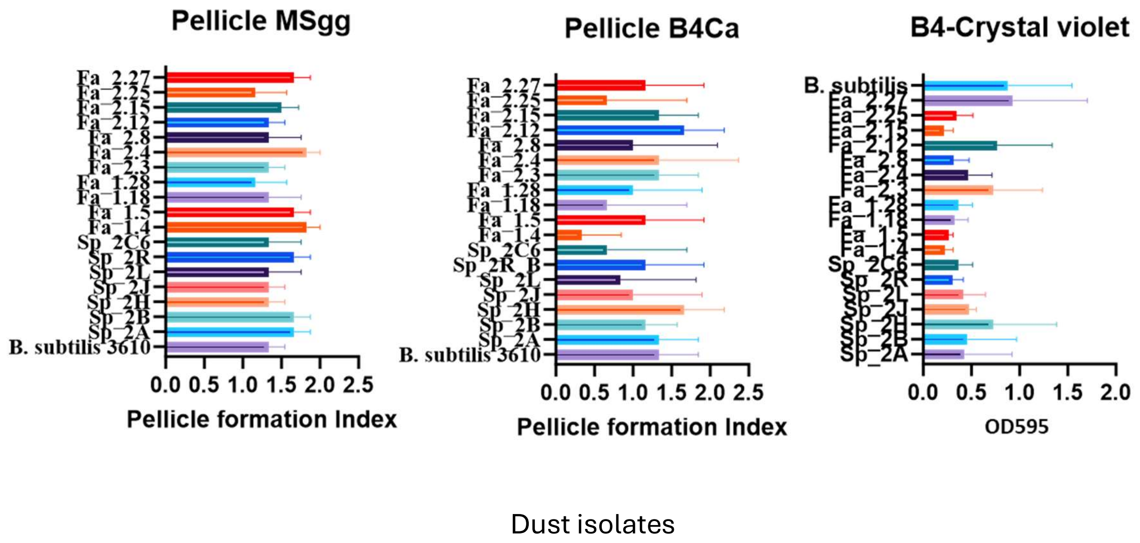


**Figure S3.** Following incubation at indicated media, pellicles were developed, and pictures were taken using Nikon Camera. Following 3 days, the experiment was repeated and insulates were grown from the same Petry dishes (that were kept at RT). Isolates were regrown by taking a colony into 2mL of LB. The Insulates were grown for ~5 hours and seeded in duplicates in either B4/B4+Ca or MSgg medium (six 24-well-plates in total) for two days incubation. In B4, biofilms were adhered to the surface and crystal violet was performed. Experiments were performed 3 independent times.

**Figure S4**

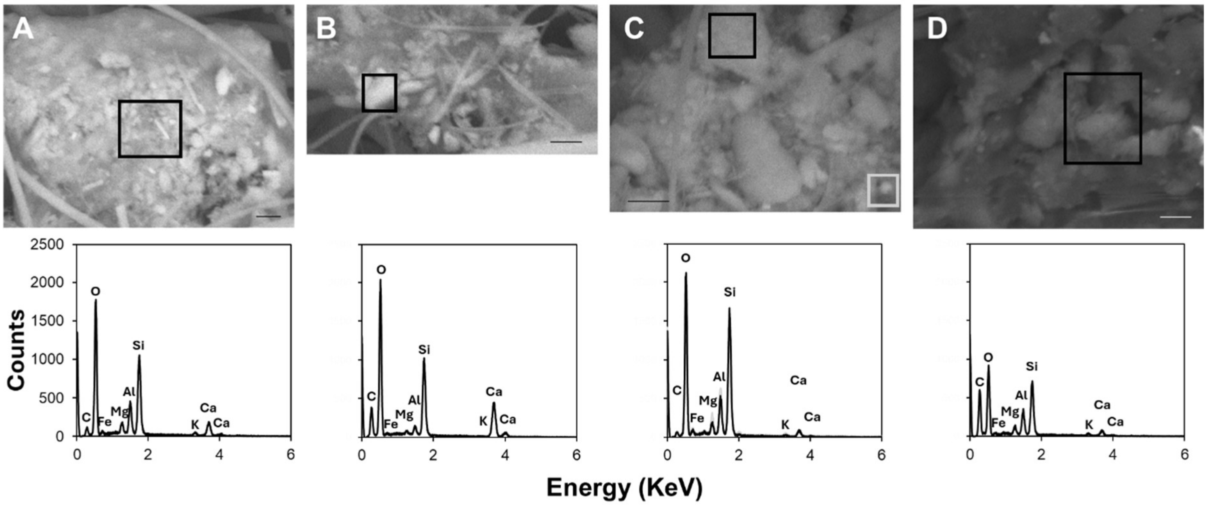


**Figure S4. Elemental composition of dust samples.** Scanning electron microscopy coupled with energy-dispersive X-ray spectroscopy (SEM-EDS) analysis was conducted for four dust particles (**A-D**) for elemental composition. Scale bars represent length of 1 µm. This representative field represent 3 independent experiments.

**Figure S5**

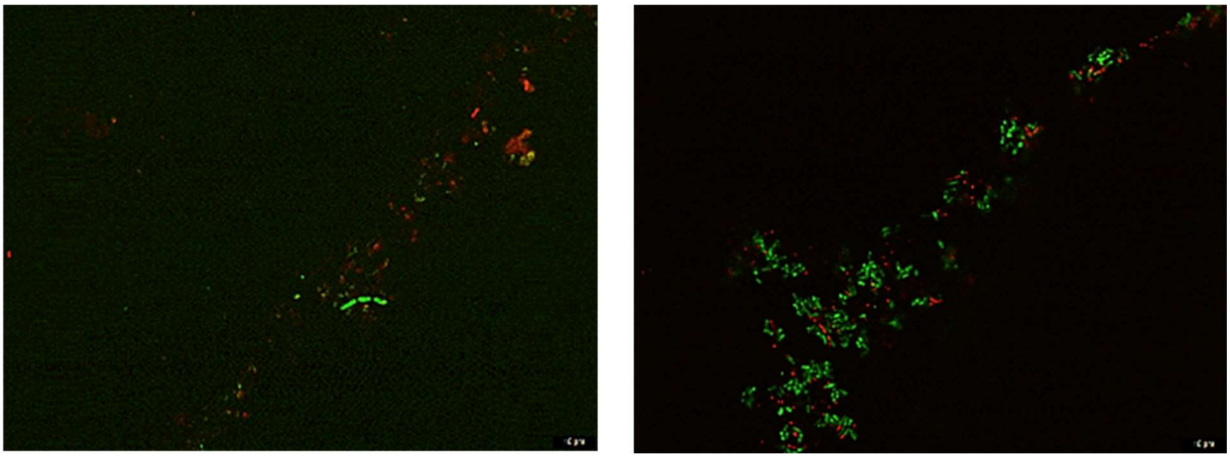

**Figure S5. Dust isolate Is_2.4 grown over sterile dust.** Growth was conducted with (right) and without (left) the addition of glutamine (Gln) as a nitrogen source and stained with a live/dead stain as described in materials and methods at indicated conditions.
